# The highly conserved RNA-binding specificity of nucleocapsid protein facilitates the identification of drugs with broad anti-coronavirus activity

**DOI:** 10.1101/2022.07.26.501505

**Authors:** Shaorong Fan, Wenju Sun, Ligang Fan, Nan Wu, Wei Sun, Haiqian Ma, Siyuan Chen, Zitong Li, Yu Li, Jilin Zhang, Jian Yan

## Abstract

The binding of SARS-CoV-2 nucleocapsid (N) protein to both the 5′- and 3′-ends of genomic RNA has different implications arising from its binding to the central region during virion assembly. However, the mechanism underlying selective binding remains unknown. Herein, we performed the high-throughput RNA-SELEX (HTR-SELEX) to determine the RNA-binding specificity of the N proteins of various SARS-CoV-2 variants as well as other β-coronaviruses and showed that N proteins could bind two unrelated sequences, both of which were highly conserved across all variants and species. Interestingly, both these sequence motifs are virtually absent from the human transcriptome; however, they exhibit a highly enriched, mutually complementary distribution in the coronavirus genome, highlighting their varied functions in genome packaging. Our results provide mechanistic insights into viral genome packaging, thereby increasing the feasibility of developing drugs with broad-spectrum anti-coronavirus activity by targeting RNA binding by N proteins.

## Main text

Following the first case reported in Wuhan, China, toward the end of 2019, COVID-19 was quickly recognized as a pandemic and has since infected more than 550 million people worldwide, claiming more than 6 million human lives as of July 2022 (according to WHO https://covid19.who.int/). COVID-19 is caused by the novel coronavirus SARS-CoV-2, which belongs to the same β-coronavirus family as the deadly SARS-CoV and MERS-CoV that caused local outbreaks in 2003 and 2012, respectively. The SARS-CoV-2 genome comprises a 29,903-nt-long positive-sense, single-stranded RNA coding for 29 viral proteins that is flanked by noncoding RNA elements, beginning with a 265-nt-long 5′-untranslated region (UTR) and a few intergenic regions and ending with a 337-nt-long 3′ poly (A) tail^1^. The 29 viral genes code for four essential structural proteins, namely the spike (S), envelope (E), membrane (M), and nucleocapsid (N) proteins, as well as 25 nonstructural accessory proteins (NSP) that have not yet been fully characterized^2^.

Upon entry into cells, SARS-CoV-2 usually hijacks cellular proteins for genome replication, protein synthesis and modifications, and virion particle packaging and release^3,4^. To understand how viral proteins selectively bind viral RNAs, we conducted high-throughput systematic evolution of ligands by exponential enrichment of RNA (HTR-SELEX), which is used to profile the sequence-binding specificity of RNA-binding proteins^5^, for all the viral proteins of SARS-CoV-2. Among all the viral proteins, strong and reproducible RNA sequence specificity was only observed for the nucleocapsid (N) protein, the most abundant protein in SARS-CoV-2 and a well-established nucleoprotein. Surprisingly, we identified two irrelevant RNA sequence motifs that were equivalently favored by the N protein; their consensus sequences were as follows: “UCCGCUUGGCC” (hereafter, motif 1) and “UAAUAGCCGAC” (hereafter, motif 2) (**Fig. 1a**; **Table S1**). Both these motifs were highly enriched in all 12 replicative experiments performed by independent researchers using different batches of SELEX-input ligands and recombinant proteins (**Extended Data Fig. 1**).

**Figure 1.**
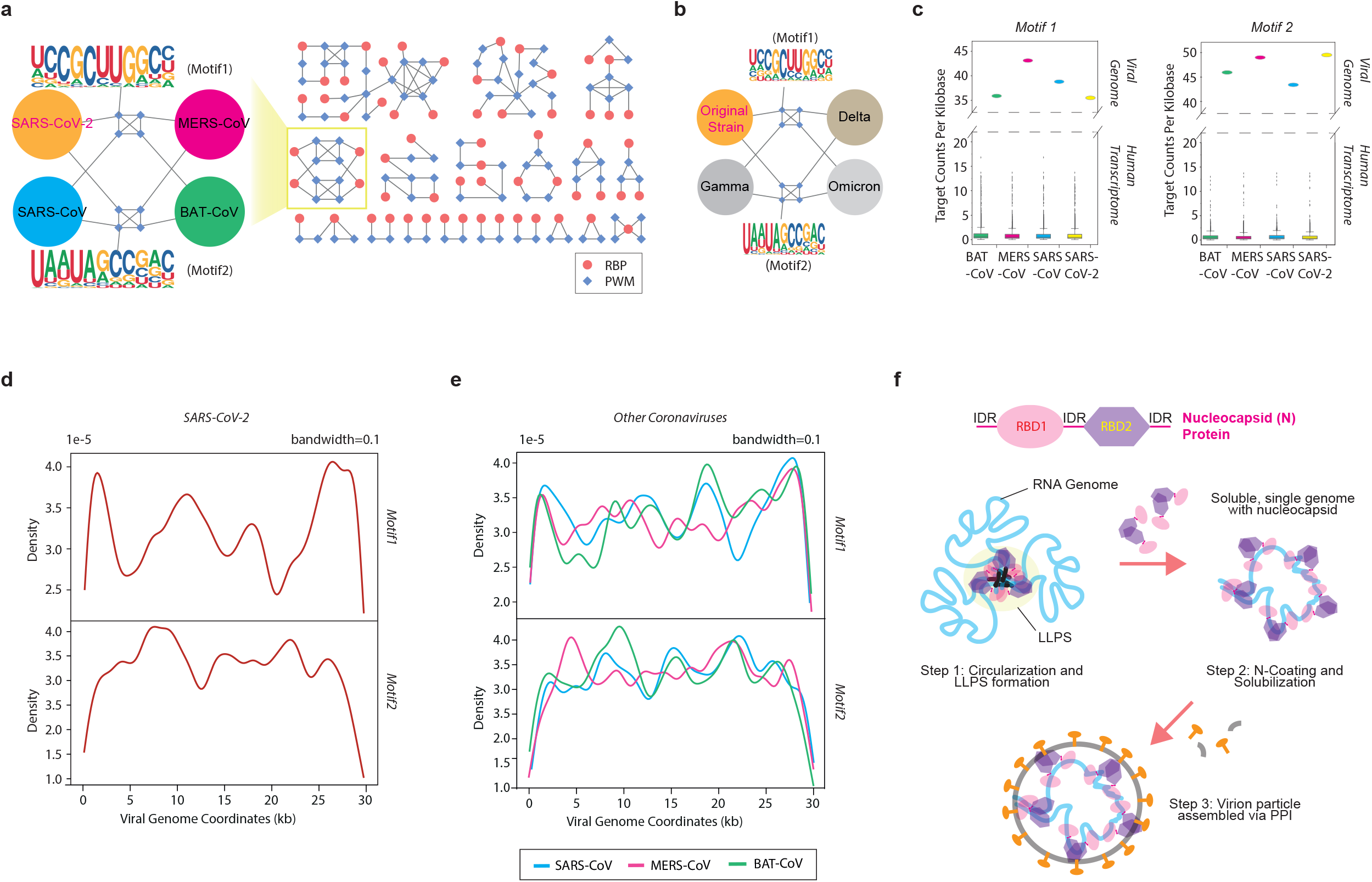
RNA binding specificity of beta-coronavirus. (a) Network analysis revealing the similarity of RNA binding specificity of RNA binding proteins, including all human RBPs with available motifs and nucleocapsid proteins of different beta-coronaviruses. Note that the subnetwork of SARS-CoV-2 nucleocapsid proteins is disconnected from other subnetworks and zoomed in for clarification. Diamonds indicate individual motifs, and circles denote RBPs. The RBP is connected to its own binding motif. The edge between motifs is drawn if the SSTAT similarity score > 1.0E-5. For details, please see **Extended Data Fig. 3**. (b) Network analysis of the similarity of RNA binding specificity of N proteins in different variants of SARS-CoV-2. The N protein of different viral variants is connected to its own binding motif. The edge between motifs is drawn if the SSTAT similarity score > 1.0E-5. (c) The comparison of binding site density of N protein on human transcriptome and viral genome. Left, binding density of motif 1. Right, binding density of motif 2. The ellipse illustrates binding density on respective viral genome, and the box plot represents binding density on human transcriptome. Green, BAT-COV; Red, MERS-COV; Blue, SARS-COV; Yellow, SARA-COV-2. (d) The binding site density (KDE, bandwidth=0.1) of N protein along SARS-CoV-2 genome. Upper, motif 1. Lower, motif 2. A binding site is significantly detected when the sequence matches the motif (P < 0.05). (e) The binding site density (KDE, bandwidth=0.1) of N protein along the genomes of the other coronaviruses. Upper, motif 1. Lower, motif 2. A binding site is significantly detected when the sequence matches the motif (P < 0.05). Smoothened color curves represent different viruses. (f) The cartoon illustrates the viral genome packaging and viron particle assembly. Upper, the structure of N protein, which includes 5’-arm, RNA binding domain (RBD1), intrinsically disordered linker region (IRD), dimerization domain (RBD2) and 3’arm. Lower, model of the viral genome packaging and viron particle assembly. In step 1, N protein forms homodimer and bind to 5’ and 3’UTRs. IDRs between different N proteins promote formation of LLPS. Subsequently in step 2, viral RNA full of motif 2 is exposed to more N proteins. Even binding of N proteins to motif 2 increases the solubility that prevents intermingling between large RNA molecules and dissolves the condensates. Finally, in step 3, the dissolved LLPS releases the packaged single RNA genome and makes the N protein accessible to interact with other viral proteins, such as M and E proteins, to complete the virion assembly.

Since the first outbreak in late 2019, multiple SARS-CoV-2 variants have emerged and have successively become dominant. The efficiency of genome packaging has been reported to be critical to the viral life cycle^6^. Therefore, we speculated whether the N protein mutations acquired by new variants promoted viral replication by enhancing the efficiency of RNA binding and virion assembly. To address this speculation, we analyzed the binding specificities of N proteins derived from three predominant SARS-CoV-2 variants: gamma, delta, and omicron (**Table S1**; **Extended Data Fig. 2a**). Interestingly, we observed no overt changes in the RNA sequence specificity of the N proteins of any of these variants (**Fig. 1b**), suggesting that the overwhelming spread of new variants was not caused by alterations in the RNA binding by N proteins but more likely due to the enhanced recognition of S proteins by cellular receptors^7^.

Besides SARS-CoV-2, multiple β-coronaviruses have demonstrated pathogenic characteristics, causing severe respiratory syndromes. For example, SARS-CoV and MERS-CoV triggered deadly epidemics and garnered global attention in 2003 and 2012, respectively. To understand the evolutionary relationships among these viruses and compare their common and differential characteristics, we performed high-throughput RNA-SELEX (HTR-SELEX), a highly sensitive method that helps detect minor differences in nucleic acid-binding specificity^8^, to examine the differential RNA-binding specificity of N proteins of these viruses (**Extended Data Fig. 2b**). We also analyzed the bat coronavirus RaTG13 isolated from horseshoe bats (from a remote cave in Yunnan Province, BAT-CoV hereafter), so far the most closely related species (sequence similarity, 96%) to SARS-CoV-2 but unable to infect humans^9,10^. Each experiment was conducted with at least four independent replicates (**Extended Data Fig. 2c**). Surprisingly, the N proteins from all these coronaviruses consistently displayed the same dual binding specificity to RNA sequences, identical to that observed for SARS-CoV-2 (**Fig. 1a**), indicating the highly conserved RNA-binding mode of N proteins as well as the consistent mode of virion packaging.

Furthermore, we compared the two motifs in terms of sequence specificity to human RNA-binding proteins (RBP) profiled using HTR-SELEX^5^; however, no significant similarity was observed with any of the known motifs (**Fig. 1a**; **Extended Data Fig. 3**). We inferred that the highly unique binding specificity of the N protein could facilitate the rapid recognition of viral RNA separated from the extremely complex pool of host cellular RNAs, even though both transcript types were generated from the host cell machinery. Such characteristics of the N protein are vital for efficient viral replication. To test this hypothesis, we scanned both the human cellular transcriptome and the four viral genomes against the two motifs. As expected, binding sites matching either of the motifs were virtually absent from the human transcriptome, with a few binding sites occasionally present in some human RNA types (**Table S2**). In contrast, hundreds of strong binding sites were significantly detected for both motifs in all the four viral genomes, consistent with the fact that approximately 720–2,200 nucleocapsid protein molecules are associated with one copy of viral RNA genome per virion^11-13^. In comparison, the density of the binding sites––both motif 1 and motif 2––is approximately 35–50 times higher in the viral genome than in the human transcriptome (**Fig. 1c**). Our results also suggested that binding of N protein to both motifs is equally important for the life cycle of the virus. It is noteworthy that the case fatality rate of β-coronavirus is somehow associated with the density of motif 1 sites in its genome. MERS-CoV—infection with which has the highest case fatality rate among all coronaviruses—encompasses the densest motif 1-binding sites in its genome, followed by SARS-CoV and then SARS-CoV-2 (**Fig. 1c**).

Next, we probed how the binding specificity of N proteins contributes to viral genome assembly and virion packaging. We scanned SARS-CoV-2 genomic RNA against each motif and constructed a density plot of the matched binding sites along the genome. Interestingly, the two motifs exhibited a mutually complementary binding pattern, i.e., the highest density of motif 1 was present in the 5′- and 3′-UTRs of the RNA genome. The relative frequency of motif 2 was low; intriguingly, motif 2, and not motif 1, was constantly enriched in the central region, ranging from approximately the 5^th^ to 25^th^ kilobases (**Fig. 1d**). The differential distribution of the two motifs underscores the potentially different roles of the UTRs and the central region in viral genome packaging. Studies have suggested that phase separation is involved in the selecting the single RNA genome and ensuring its compactness^14-16^. In such a model, the binding of N proteins to the 5′- and 3′-UTRs promotes liquid–liquid phase separation (LLPS), whereas the association of N proteins with the central region enhances the fluidity and solubilization of the viral genome and limits its entanglement with other large RNA molecules, thereby increasing the packaging efficiency^17,18^. In coronaviruses in particular, when the viral genome is packaged, the association of N proteins with the central regions dissolves the condensates and anchors the single RNA genome onto the viral membrane via interactions between the N protein and the M protein^14,19^.

Although the RNA-binding specificity of N proteins is highly conserved among coronaviruses, the genomic sequence composition could be highly diverse. Thus, coronaviruses could still undergo genome packaging via different fashions. For comparison, we mapped the N protein-binding site densities along individual genomes and noted that the distributions of both binding site types were highly similar to those of SARS-CoV-2 (**Fig. 1e**), confirming that β-coronaviruses go through a conserved packaging process mediated by the dual binding of N proteins with different viral genome regions.

The structure of the SARS-CoV-2 N protein has been successfully determined; it contains two highly structured domains separated by three intrinsically disordered regions (IDRs), an N-terminal RNA-binding domain (RBD1), and a C-terminal dimerization domain that also possesses RNA-binding ability (RBD2)^20-23^ (**Fig. 1f**). We suspected that RBD1 and RBD2 were responsible for recognition of motifs 1 and 2, respectively. Therefore, we individually cloned the two domains into an *E. coli* expression vector and isolated soluble proteins for HTR-SELEX analysis (**Table S1**; **Extended Data Fig. 4**). Each experiment was independently conducted at least thrice. We barely observed any enrichment of either motif in any of the replicative experiments (data not shown), which indicates that RBD1 or RBD2 alone cannot stably bind any specific RNA sequence. Given the role of RBD2 in homodimerization of N protein, our results suggest that N protein dimerization is also required for its binding to either motif. This difference in the sequence specificity can be attributed to certain topologies of the N protein homodimers. To this end, we propose a model stating that during virion particle replication initiation, N protein dimers bind the 5′- and 3′-UTRs, provoking LLPS via IDR aggregation (linker region) and consequently circularizing the RNA genome. Such a change results in the exposure of the central region for subsequent coating with more nucleocapsid proteins. Because motif 2 was highly enriched, the binding of N proteins to such sites in the central region mostly displayed a different topology compared with that in the UTRs and hidden in IDRs. The binding of N proteins to the central region increased the fluidity and solubility of the N protein–RNA complex and ultimately allowed its interaction with other structural and accessory components, consequently leading to virion particle assembly (**Fig. 1f**).

To control the severe health and economic impacts of COVID-19, extensive effort has been made to identify effective drugs that can prevent its spread and ease its symptoms. Many antiviral drugs, including monoclonal antibodies, target the S protein. Owing to the fast mutation rate of SARS-CoV-2, the emergence of new variants can quickly mitigate drug efficacy^24,25^. Thus, it is more reasonable to target the less-mutable N protein. Small chemical compounds or synthesized peptides that are cytoplasmic membrane-permeable have been used to inhibit dimerization or RNA binding or to dissolve the LLPS in antiviral treatment^26-28^. For example, PJ-34 and H3, which are small-compound inhibitors of the N-terminal RNA-binding domains of N proteins, were found to be effective in impeding the replication of the human β-coronavirus HCoV-OC43^29^. During the MERS-CoV outbreak, 5-benzyloxygramine (P3) was identified to inhibit N proteins by promoting abnormal aggregation. At concentrations of 100 µM, 5-benzyloxygramine (P3) could completely prevent viral production and replication^30^. Based on the high binding specificity of the N protein to the viral genome but not the host transcriptome, we could synthesize and deliver, as antiviral drugs, short-sequence single-stranded RNA to cells, with the sequence either identical or complementary to the consensus sequence “UCCGCUUGGCC.” Such small RNA molecules can prevent viral replication by competing the binding between the viral genome and N protein. Given that both the RNA binding and genome composition are highly conserved among all β-coronaviruses, antiviral drugs targeting the RNA binding of N proteins could help fight against a broad spectrum of β-coronaviruses and can therefore be used in treating any novel coronavirus-related diseases that may occur in the future. Similar studies to assess the RNA-binding specificity of N proteins among other viral species (e.g., HIV, HBV, and Ebola virus) will continue, and the findings of the current and future studies will gradually expand our understanding of the process of viral genome packaging and facilitate the identification of effective therapies.

## Online Methods

### Vector construction and protein purification

Different coronavirus N proteins and the RNA Binding Domains (RBDs) of SARS-CoV-2 were cloned into pET28a expression vector with an N-terminal 6×His tag using the recombinase (Vazyme ClonExpress II One Step Cloning Kit, Vazyme Biotech). The resulting plasmids were transformed into *E*.*coli* Rosetta (DE3) strain for protein purification. Then, the bacterial was cultured in LB and induced with 0.1 mM IPTG when OD_600_ reached 0.6, continued to be cultured for 16 h at 16 ºCwith rigorous shaking. The bacterial culture was centrifuged 13000 ×g for 5 min, and the pellet was resuspended in Buffer A (0.5 mg/ml Lysozyme, 50 mM tris-HCl pH 7.5, 300 mM NaCl, 10 mM Imidazole, 1 mM DTT, 1 mM PMSF) at 4 °C, standing for 1h and ultra-sonicated for 10 min at an interval of 10 seconds, followed by centrifugation at 13,000 × g for 30 min at 4 °C. The supernatant was loaded into a Ni-NTA column (Bio-Rad). The Ni-NTA beads were set at 4 °C for 30 min before being eluted with Buffer A containing 300 mM imidazole. Finally, the eluted proteins were analyzed with SDS-PAGE to confirm its purity and molecular weight. The cDNA and amino acid sequences of all protein used in this study could be found in **Supplementary Table S1**.

### HTR-SELEX

The HTR-SELEX experimental procedure was modified from the previous study^5^. Briefly, we first produced the DNA templates of HTR-SELEX containing a T7 promoter by PCR amplification. RNA input was transcribed from the DNA-templates using T7 *in vitro* transcription (HiScribe™ T7 Quick High Yield RNA Synthesis Kit, NEB, E2050S) following the manufacture’s instruction. The remaining DNA was digested by DNaseI (RNase free). To enable the RNA to form secondary structures, the RNA input libraries were then heated to 70°C followed by slow cooling. After that, 600 ng of N protein was mixed with 5 µL of RNA ligands (SELEX input) in 20 µL binding buffer (50 mM NaCl, 1 mM MgCl_2_, 0.5 mM Na_2_EDTA, 1U/µL RNase inhibitor and 4% glycerol in 50 mM Tris-Cl, pH 7.5). After incubated for 15 minutes at 37 °C, the mixture was continued to be incubated at room temperature for 15 min. Subsequently, 150 µL binding buffer which contained 10 µL of pre-equilibrated Ni Sepharose 6 Fast Flow resin (GE Healthcare, 17-5318-01) was added into the protein-RNA mixture, followed by 2 hours gentle shaking at room temperature. The beads were then repeatedly washed with 200 µL of binding buffer 12 times to remove the unbound RNA ligands. After removing the supernatant by centrifugation at 1000 ×g, the beads were resuspended in 20 µL of elution buffer (0.5 µM RT-primer, 1 mM EDTA and 0.1% Tween 20 in 10 mM Tris-Cl buffer, pH 7.0) and heated to 70°C for 5 minutes followed by cooling to 4 °C to allow the reverse transcription primers annealed to the RNA. Finally, reverse transcription was conducted followed by PCR amplification using the barcoded amplification primers listed in **Supplementary Table S1**. The obtained PCR products were used as DNA templates for the next HTR-SELEX cycle. This SELEX process was repeated four times. PCR products from each SELEX cycle were also purified and sequenced using BGI MGISEQ 2000 or illumina Novaseq sequencer.

### HTR-SELEX data analysis

The data were binned according to barcodes for each sample. After discarding the low-quality reads, the remaining sequences were trimmed to remove adaptor sequences. The remaining 40-nt region were subjected to further analysis. The position weight matrix (PWM) was generated as described in Jolma et al^5^. Briefly, the in-house software Autoseed identified the most frequent gapped and un-gapped k-mer sequences representing local maximum counts relative to similar sequences within the Huddinge neighborhoods^31^. Count of each seed motif is higher than that of any similar sequence within a Hamming distance of one. The initial set of motifs is manually curated to make sure that the final seeds did not include partial motifs that included constant linker sequences (displayed a strong positional bias on the ligand), and motifs that were recovered from a large number of experiments. PWM models were built from sequences matching the indicated seeds, allowing sequencing reads only up to Hamming distance of 1. Individual results that were not supported by replicate were deemed inconclusive and were not included in the final dataset. Draft models were manually curated (by JY, LF and SF) to remove unsuccessful experiments and artefacts due to bottlenecks and aptamer selection. After that, the uracil was replaced with thymine to adapt RNA analysis. Then, we used the R package ggseqlogo to construct seqlogos for all PWMs. Final models for each replicative experiment were generated and stored in **Table S1**.

### Network analysis of similarity between PWMs

The SSTAT (parameters: 50% GC-content, pseudocount regularization, type I threshold 0.01) was used for calculated the similarities of human RBPs and coronavirus N protein PWMs as described in Fan et al^32^. After that, we construct a similarity network in which diamonds indicate motifs and circles indicate RBPs. The RBPs were connected to their motifs, and the different motif were further connected if the SSTAT similarity score > 1.5E-5. Cytoscape software (v3.7.2) was used for visualizing the network.

### Binding site density analysis

The genome-wide target sites were scanned using FIMO (v5.0.4, Grant CE, bioinformatics, 2011) for all motifs. All predicted binding sites are available in **Table S2**. To construct **Fig 1c**, binding site density was transformed to Target Counts Per Kilobase by using a customer script (available upon request). The kdeplot fuction of python package seaborn was used for calculate and plot binding site density under parameter bandwidth=0.1.

## Acknowledgments

This work was financially supported by the National Natural Science Foundation of China (81873642, 32070596, 31900443), the Research Grants Council of Hong Kong SAR (21100420, 11101022), City University of Hong Kong (9667240, 9610424, 7005747), the Tung Biomedical Sciences Centre (9609303), and Hetao Shenzhen-Hong Kong Science and Technology Innovation Cooperation Zone Shenzhen Park Project (HZQB-KCZYZ-2021017).

## Author contributions

J.Y., J.Z., S.F., L.F. and W.J.S. conceived the project. S.F., L.F., and N.W. carried out experiments, J.Z., W.J.S., W.S., Z.L., S.C., and Y.L. performed data analysis. J.Y., J.Z., L.F., S.F., and W.J.S. wrote the manuscript.

### Data and Code Availability

Sequencing data generated in this study can be accessed via Gene Expression Omnibus (GEO). All primer sequences used in this study are available in **Table S1**.

**Extended Data Fig. 1.**
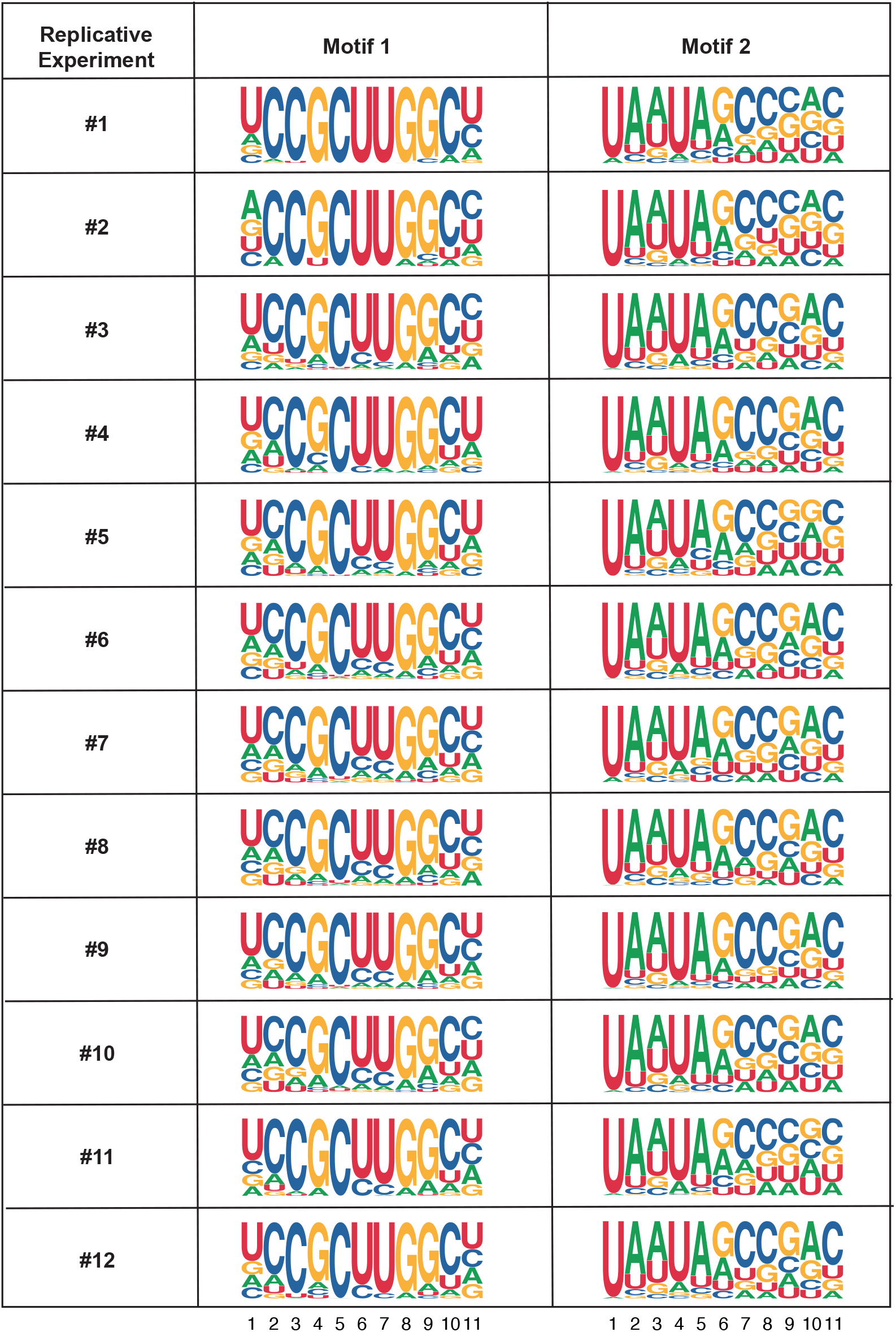
Binding motifs of N proteins of SARS-CoV-2 with 12 replicative experiments. HTR-SELEX results reveal that the two motifs are highly enriched in all 12 replicative experiments.

**Extended Data Fig. 2.**
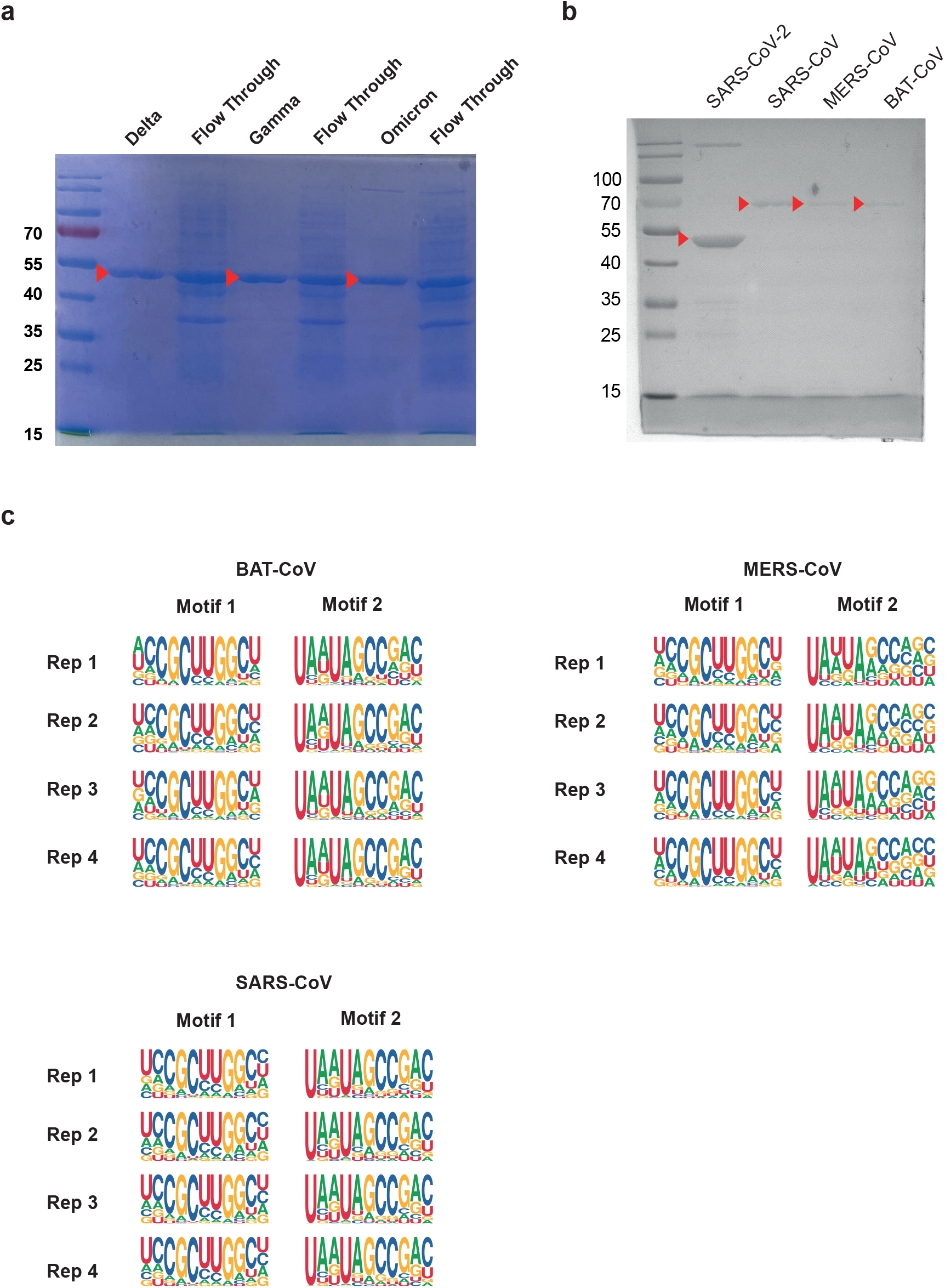
Expression of recombinant N proteins of different viral species with *E*.*coli*. (a) The SDS-PAGE gel, stained with Coomassie brilliant blue, showed the expression of N proteins from three SARS-CoV-2 variant strains including the Flow Through next to it. (b) The SDS-PAGE gel, stained with Coomassie brilliant blue, showed the expression of N proteins from different coronaviruses. (c) RNA binding specificity of N proteins from different coronaviruses, each with 4 independent replicates

**Extended Data Fig. 3.**
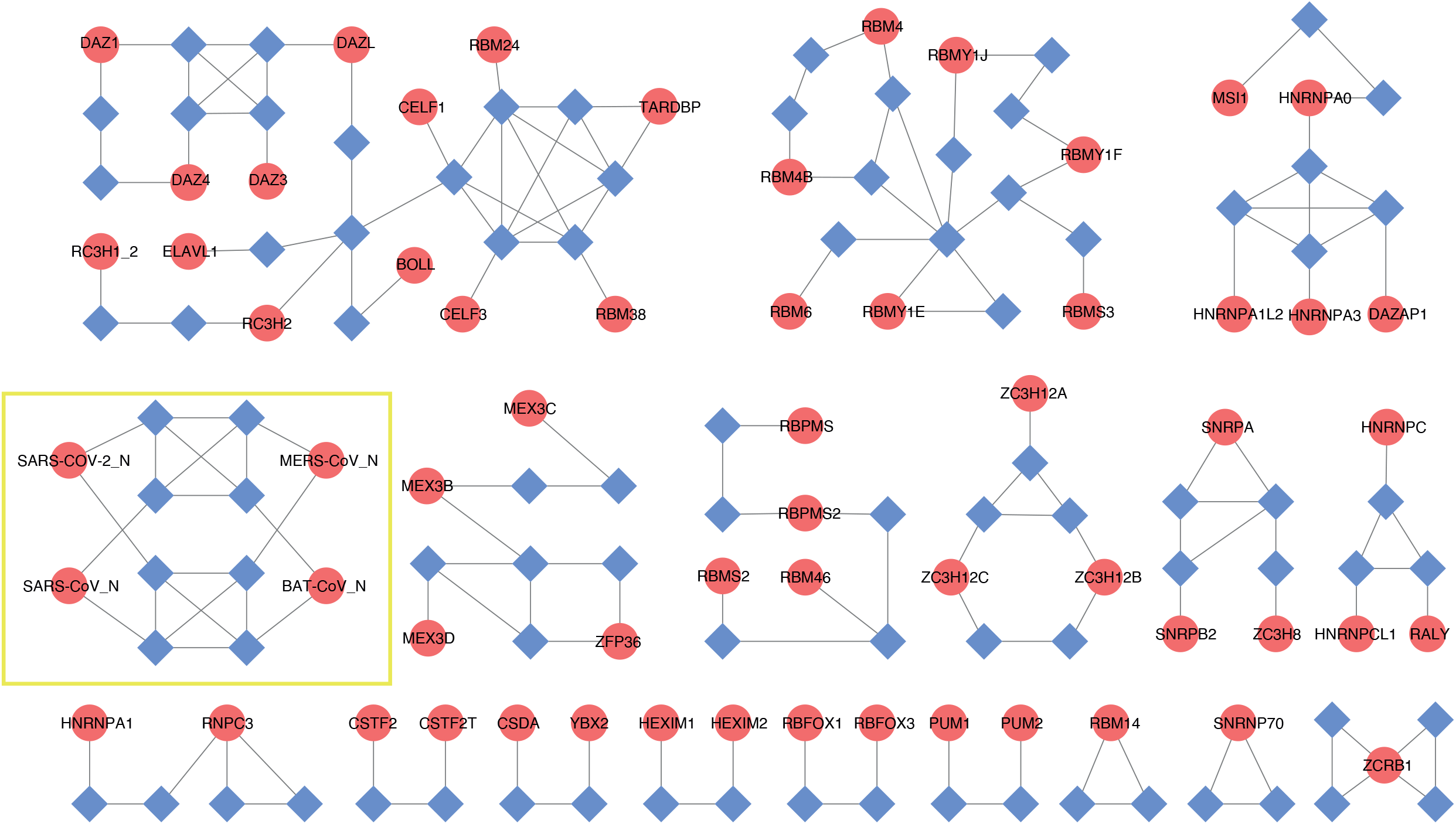
Network analysis of similarity between human RBPs and coronavirus N protein PWMs. The zoomed illustration of **Fig. 1a**. Diamonds denote motifs and circles represent RBPs. The RBPs are connected to their motifs, and the motif are further connected to each other if the SSTAT similarity score > 1.0E-5. Coronavirus N protein network are framed in yellow because it has no similarity with human RBPs.

**Extended Data Fig. 4.**
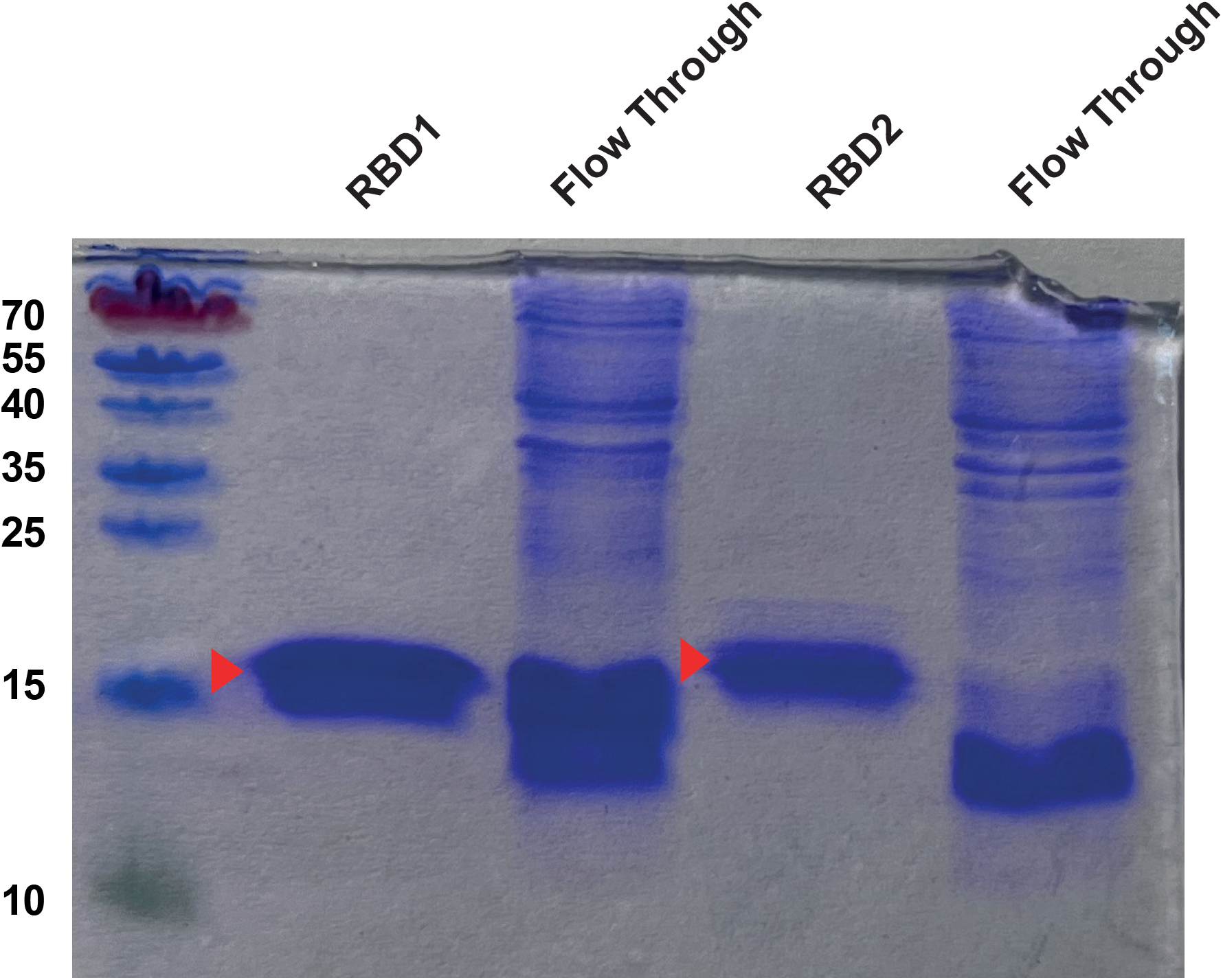
Expression of RBDs of SARS-CoV-2. The SDS-PAGE gel, stained with Coomassie brilliant blue, showed the expression of two RBDs of N proteins from SARS-CoV-2 including the Flow Through next to it.

